# PAQman: reference-free ensemble evaluation of long-read genome assemblies

**DOI:** 10.1101/2025.09.11.675652

**Authors:** Samuel O’Donnell, Ningxiao Li, Jacob L. Steenwyk, David M. Geiser, Frank N. Martin, Emile Gluck-Thaler

## Abstract

Advances in long-read sequencing have made it easier and more cost effective to generate high-quality genome assemblies. However, assessing assembly quality remains a challenge, as existing tools often focus on a few metrics and/or require a reference assembly for comparison. Furthermore, the number of available metrics and associated tools for genome evaluation have expanded in recent years, making it more difficult for researchers to easily use and develop comprehensive pipelines. To address this, we developed the Post-Assembly Quality manager (PAQman), a tool that lowers the barrier to entry for assembly quality assessment by measuring seven reference-free features of genome quality within a single framework: Contiguity, Gene content, Completeness, Accuracy, Correctness, Coverage, and Telomerality. PAQman integrates multiple commonly used tools alongside custom scripts, requiring users to provide only a query genome assembly and its underlying long-read data, while providing a streamlined and consistent framework for quality assessment across datasets.

**Impact Statement:** PAQman is an ensemble-based tool for comprehensive, reference-free evaluation of genome assemblies derived from long-read sequencing data. The simultaneous integration of seven quality features enables users to easily assess assembly quality within a standardized, reproducible framework across diverse organisms, while lowering the barrier to entry for biologists analyzing their data.

## Introduction

The ease with which researchers can generate reference-quality assemblies has advanced rapidly with the continued improvement of long-read sequencing technologies, namely those offered by Oxford Nanopore Technologies (ONT) and Pacific Biosciences. In parallel, assembly algorithms and pipelines have significantly improved in both accuracy and speed (Li 2018; Kolmogorov et al. 2019; Giani et al. 2020; Espinosa et al. 2024). Despite these advancements, assembly quality is often assessed by disparate software that individually use a subset of available metrics, leading to inefficiencies in throughput and evaluation. These different statistics often evaluate distinct genomic features and thus do not always correlate with one another (Zhang et al. 2023). Determining the most appropriate metrics for quality assessment is therefore challenging, especially when benchmarking software or parameter settings to optimize assembly quality for a particular dataset or organism (Bradnam et al. 2013; Zhang et al. 2022; Cosma et al. 2023). These difficulties are further compounded in the absence of a reference genome, where defining the “best” assembly becomes even more ambiguous.

Several tools have been developed to address the challenge of assembly quality assessment, such as Genome Assembly Evaluation Process (GAEP) (Zhang et al. 2023), GenomeQC (Manchanda et al. 2020) and the most commonly used QUAST pipeline (Mikheenko et al. 2023). However, most are either sparsely documented, challenging to install and/or use, or evaluate only a subset of relevant statistics. For example, GenomeQC, if used without a reference assembly or gene annotations, only calculates a few assembly stats such as contig N50 and gene content using BUSCO. Similarly, QUAST relies on comparisons with a reference assembly to compute several statistics, such as structural accuracy and completeness. Although GAEP evaluates a number of important statistics such as contiguity, gene content, accuracy and structural errors, it does not assess coverage, completeness or the presence of telomeric sequences, and is not packaged for use as a software container or conda package for use in high-throughput computing environments such as those commonly used for bioinformatics analyses.

To address these challenges, we developed the Post-Assembly Quality manager (PAQman), a tool that calculates seven reference-free features of genome quality—Contiguity, Gene content, Completeness, Accuracy, Correctness, Coverage and Telomerality—by integrating multiple existing and popular tools and custom scripts. PAQman requires only the assembly itself and the long-read data used to generate it, while offering users an accessible and standardized approach for genome quality assessment in high-throughput computing environments. Although PAQman was designed with eukaryotes in mind, there is no limitation in organism type aside from the telomerality feature being designed for linear chromosomes found primarily in eukaryotes.

## Methods

The workflow summarized in Fig. 1 describes the two commands used in PAQman. The first, *paqman*, performs the separate evaluation of seven features of assembly quality prior to compiling the summary statistics table. By default, the minimum requirements for this command are an assembly in FASTA format (-a) and long-reads in FASTQ format (-l). Additional short paired-end reads may also be provided (-1 -2; FASTQ). Each feature and the tool/s used for its evaluation are outlined below:

**Figure 1.**
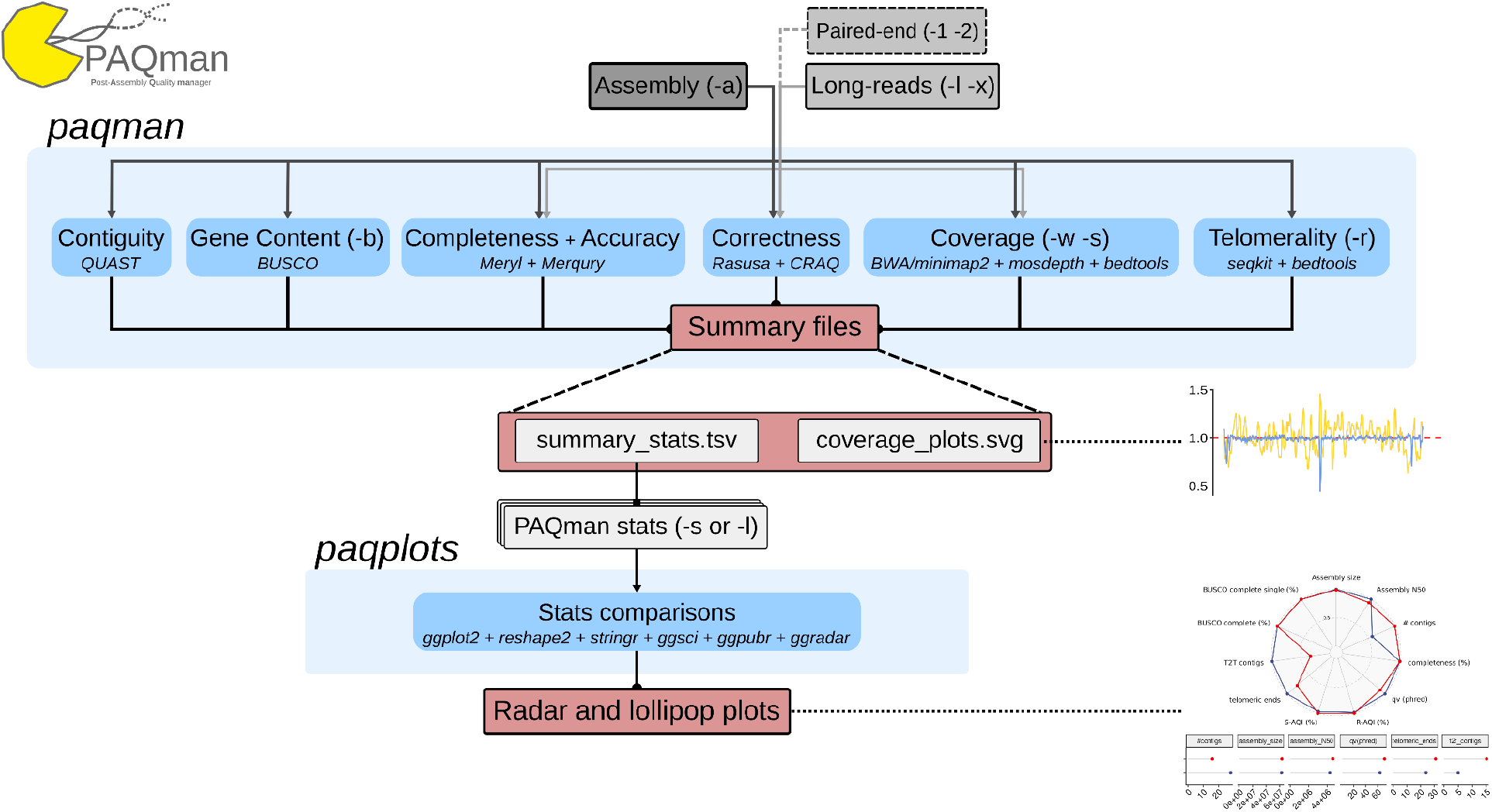
Schematic of the PAQman pipeline consisting of the *paqman* and *paqplots* commands. Grey boxes indicate files, blue boxes indicate computational steps calculating specific assembly metrics (with software names italicized below) and red boxes indicate outputs. Example figures output by PAQman are to the right. Brackets indicate the short-hand command line arguments for input files and parameters impacting that specific step.

### Feature 1: Contiguity

By far and above, the most common metrics evaluated post-assembly are the number of contigs/scaffolds, the assembly size and the contig/scaffold N50. PAQman runs QUAST (Mikheenko et al. 2023) to rapidly calculate these important metrics. Generally speaking, a lower number of scaffolds and larger assembly size and contig N50 are associated with higher quality assemblies. Additional analyses from QUAST, such as GC content and cumulative length plots, are kept in the output folder.

### Feature 2: Gene content

Apart from contiguity, the most common tool used to evaluate assemblies is BUSCO, which quantifies the recovery of various databases of universally conserved single copy orthologs (Tegenfeldt et al. 2025). Using the presence/absence of near-universal single-copy orthologs, BUSCO gives a proxy for assembly completeness in terms of gene content. PAQman parameter *-b* allows users to provide the most appropriate database for BUSCO to use given the taxonomic lineage of the focal organism (more database information at https://busco.ezlab.org/busco_userguide.html#obtain-busco). The percent recovery of single-copy orthologs (ranging from 0-100%) is positively associated with assembly quality. Users can also provide a predownloaded or locally generated BUSCO database and provide it to PAQman using *--localbuscodb*.

### Feature 3: Completeness

Genomic regions containing a high density of protein-coding genes are generally the easiest to assemble due to relatively low repetitiveness compared to non-genic regions. Therefore, in addition to gene content, assembly completeness may also be more globally evaluated in terms of whether *k*-mers present in the raw read inputs are also present in the assembly. The *k*-mer-based completeness statistic is thus an accurate representation of how much of the raw sequencing data is present in a final assembly, including both repetitive and non-repetitive content. This *k*-mer-based measure of completeness is therefore distinct from other uses of the term that are frequently used to describe assembly quality, such as the gene content-based completeness measurement implemented in BUSCO. PAQman first builds a *k*-mer distribution of the reads, selecting the short-reads (-1 -2) if provided, using meryl *count* (Miller et al. 2008) then provides this meryl database and the assembly to Merqury (Rhie et al. 2020) to calculate the *k*-mer based completeness. Completeness is calculated as a percentage with a maximum of 100%. Users may optionally provide a precomputed meryl database using *--meryldb*. A limitation of this unbiased evaluation of raw reads is that the resulting *k*-mer based metrics can be affected by contamination, mixed samples, or complex ploidy, complicating the interpretation of completeness and QV estimates.

### Feature 4: Accuracy

Using the *k*-mer distributions calculated for Feature 3, Merqury also estimates the number of genome-wide sequencing errors and provides a Phred quality score (=-10×log_10_(Pe); Pe: estimated probability of error) for the assembly. This estimate indicates how many errors may still be present in an assembly and may therefore be used to test whether common assembly polishing tools/pipelines improve quality. In general, assemblies with a Phred Quality greater than 30 are considered of high quality, although 45+ is common with more recent long-read chemistries and polishing methods. In rare cases where no errors are detected, we used the rule of three to calculate a conservative Phred score estimate, substituting the estimated probability of error by 3/*n*, where *n* is the genome size.

### Feature 5: Correctness

As genome assemblies approach reference quality, assessment must go beyond simple presence or absence of genomic content to include whether that content is assembled in the correct genomic location and structure. CRAQ (Li et al. 2023) is a tool that uses read mapping evidence to highlight regions within an assembly that have potential assembly errors. This provides an estimate for genome-wide structural accuracy, thereby providing evidence to support claims of assembly contiguity. Initially, PAQman randomly down-samples the provided long-read sequencing data to a maximum of 30X coverage by default (set by the *--maxcoverage* parameter) using Rasusa (Hall 2022), and then provides the filtered read dataset to CRAQ in addition to short-reads, if provided. PAQman then focuses on two versions of the Assembly Quality Index (AQI) summary statistic; the R-AQI and S-AQI, that score the assembly quality based on the detection of small regional (R) and large structural (S) errors respectively. The AQI scores are bound between 0-100, with 100 meaning the absence of detectable errors. Please refer to Li et al., 2023 for a complete description on how both R- and S-AQI are calculated.

### Feature 6: Coverage

Using read mapping, PAQman also evaluates whether coverage varies in particular regions compared to the genome-wide median. One important use of this evaluation metric is to determine whether multi-copy repeat regions have collapsed into a single copy during assembly, as coverage will noticeably increase in collapsed regions. Coverage may therefore help in understanding and locating more complex rearrangements such as large duplications and aneuploidies. PAQman uses bwa mem (Li 2013) and/or minimap2 (Li 2018) for short- and long-read mapping respectively; mosdepth (Pedersen and Quinlan 2018) and bedtools (Quinlan and Hall 2010) for pre- and post-processing; and custom Rscripts with ggplot2 (Wickham 2016) for plotting the relative coverage normalised by the genome-wide median (Fig. 1). PAQman uses the randomly downsampled long-reads for mapping to improve speed. The parameters *-w* and *-s* control the size of the window and slide, respectively, used to average coverage across the genome. For a final statistic, PAQman calculates the percentage of the genome that is within two standard deviations of the genome-wide median coverage. Ideally, larger percentage values are more desirable.

### Feature 7: Telomerality

This term defines a set of statistics that indicate whether an assembly contains assembled telomeric ends, which is a good indication that complex subtelomeric regions have been assembled and that contigs may thus represent full telomere-to-telomere (T2T) chromosomes. In most cases, this statistic will be used to elevate assemblies ranging from very good quality to complete. This feature can be ignored in organisms with circular DNA molecules. Organism-specific telomeric repeats are provided through the *-r* parameter, quickly identified using seqkit *locate* (Shen et al. 2016) and processed using bedtools *merge* for overlapping repeats with a max of one missing repeat between. Repeats are considered telomeric if the aggregated repeat regions are at least the size of two repeats. Contig/scaffold ends are considered to be capped by telomeres if the distance of a repeat to an end is less than 75% of the size of the entire telomeric repeat region. We recommend using TeloBase (Lyčka et al. 2024) in order to find the likely telomeric sequence for your species, however this database does not allow for degenerate repeats. To help with this, we are compiling a list (https://github.com/SAMtoBAM/PAQman/blob/main/telomeric_sequences.md) of working telomeric repeat patterns (both exact and regular expressions) found to work in manually verified assemblies.

The second optional command in PAQman, *paqplot*, takes the summary statistics output by *paqplot* from multiple assemblies and generates radar and lollipop plots to help benchmark multi-assembly comparisons using both the raw and relative values (Fig. 2).

**Figure 2.**
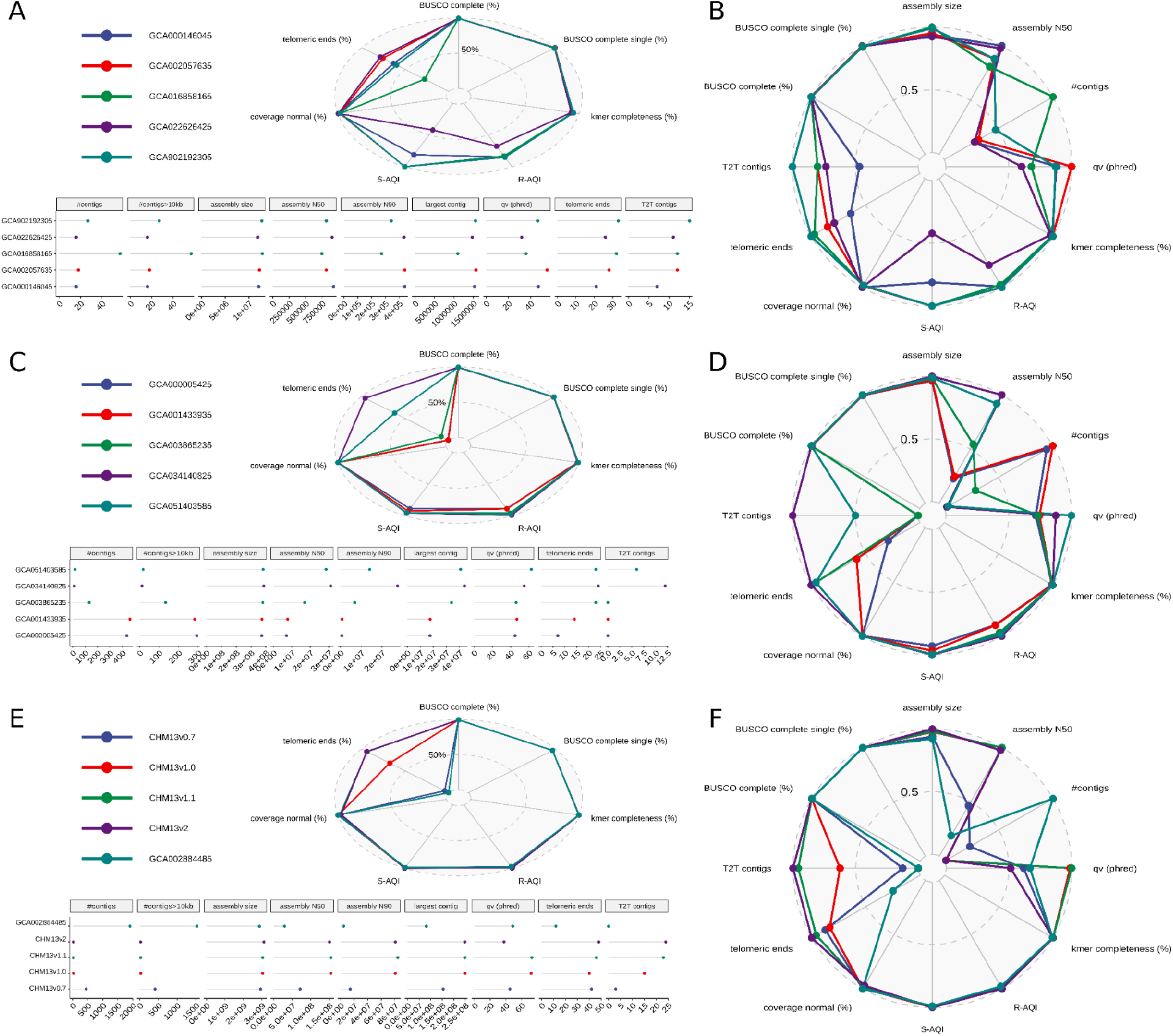
Examples of output visualizations from paqplot outputs of various paqman Feature metrics. Three different datasets were used: (A-B) *S. cerevisiae* reference strain S288c, (C-D) *Oryza sativa* subsp. *japonica* Nipponbare cultivar, and the human cell line CHM13; each with five different assemblies. (A, C, E) Radar and lollipop plots using the raw values of several metrics. (B, D, F) Radar plot using relative scales of metrics between assemblies. The individual metrics plotted represent all seven Features of assembly quality measured by PAQman; 1. Contiguity (‘assembly size’, ‘assembly N50’, ‘assembly N90’, ‘largest contig’, ‘#contigs’ and ‘#contigs>10kb’); 2. Gene content (‘BUSCO complete single (%)’ and ‘BUSCO complete (%)’); 3. Completeness (‘kmer completeness (%)’); 4. Accuracy (‘qv (phred)’); 5. Correctness (‘R-AQI (%)’ and ‘S-AQI (%)’); 6. Coverage (‘coverage normal (%)’); and 7. Telomerality (‘Telomeric ends (%)’, ‘Telomeric ends’ and ‘T2T contigs’).

## Results and Discussion

We tested PAQman on three different genome datasets with diverse genome characteristics: *S. cerevisiae* (S288c), *Oryza sativa subsp. japonica* (Nipponbare) and Humans (CHM13); encompassing a range of eukaryotic lineages (fungi, plants, and animals), genome sizes (∼12Mbp, ∼380Mbp, and ∼3Gbp), repeat content (∼10%, ∼40%, and ∼55%) and ploidy (haploid, diploid, and effectively haploid). Notably, each dataset contains multiple independent and/or sequentially improved versions of the strain/cultivar/line with an ultimately T2T version. For each dataset (S288c, Nipponbare and CHM13), a complete, reproducible and detailed walkthrough can be found here https://github.com/SAMtoBAM/PAQman/tree/main/pub_examples/.

For the *Saccharomyces cerevisiae* reference strain S288c, we tested PAQman on a dataset of five publicly available assemblies (GCA_000146045.2, GCA_902192305.1; GCA_022626425.2; GCA_002057635.1; GCA_016858165.1). Each was evaluated using the same ONT dataset (SRR17374240) (Zhang et al. 2022) downsampled from 800x to 100x coverage using Rasusa (*-b 1200000000*). *paqman.sh* was run with default options except ‘*-b saccharomycetaceae*’ and ‘*-r GGTGTG*’ for the BUSCO db and telomeric repeat, respectively. All summary stats were then evaluated by *paqplots.sh* using default settings (Fig. 2A-B). PAQman shows that although assembly GCA022626425 (purple) contains good metrics for the two most commonly evaluated features, Contiguity (fewest contigs and high contig N50) and Gene Content (high single copy BUSCO content), it contains poorer metrics for other less commonly used features such as Accuracy and Correctness. The lower AQI values in GCA022626425, indicative of assembly errors, were confirmed by whole genome alignment (data not shown). Similarly, GCA000146045 (teal), the canonical reference assembly, contains good relative metrics for all features except Telomerality, with only 21 contig ends identified as containing telomeric repeats, the same as previously detected (O’Donnell et al. 2023). These two examples highlight how each Feature is an independent dimension and that only a comprehensive evaluation of all features allow us to accurately measure quality.

For the *Oryza sativa subsp. japonica* Nipponbare cultivar, we ran PAQman on five publicly available assemblies (GCA_003865235.1, GCA_051403585.1, GCA_000005425.2, GCA_001433935.1, GCA_034140825.1) which includes the previous reference assembly (GCA_001433935.1) and the new gold standard T2T assembly (GCA_034140825.1). We analysed all assemblies using PacBio HiFi (SRR25241090) reads (∼85X) from the T2T project (Shang et al. 2023). *paqman.sh* was run with default options except ‘*-b poales*’ and ‘*-r TTTAGGG*’ for the BUSCO db and telomeric repeat respectively. All summary stats were then evaluated by *paqplots.sh* using default settings (Fig. 2C-D). The results show that the assemblies GCA051403585 (teal) and GCA034140825 (purple) have the best set of metrics for all Features, including Accuracy (‘qv-phred’). These two assemblies only differentiate largely due to Telomerality, as the reportedly T2T assembly (GCA034140825; purple) indeed contains a full set of 12 chromosomes all capped by telomeres (‘T2T contigs’) (Shang et al. 2023). This difference highlights the benefit of Telomerality as a feature of assembly quality.

For the human cell line CHM13, we ran PAQMan on five publicly available assemblies (T2T-CHM13v0.7, T2T-CHM13v1.0, T2T-CHM13v1.1, T2T-CHM13v2.0 and GCA_002884485.1). This includes 4 versions of the now telomere-2-telomere, complete assembly (T2T-CHM13v2.0) and one much earlier assembly for the same cell line (GCA_002884485.1). We also used PacBio HiFi (SRX5633451) reads from the T2T-CHM13 project (https://github.com/marbl/CHM13) with approximately 24X coverage in total. *paqman.sh* was run with default options except ‘*-b tetrapoda*’ for the BUSCO db and *--coveragemax 0* to skip downsampling due to low input coverage. All summary stats were then evaluated by *paqplots.sh* using default settings (Fig. 2E-F). As expected we see that the Contiguity and Telomerality features improve with subsequent versions of the T2T CHM13 versions and that v0.7 already drastically improved upon the previous public accession (GCA002884485). This once again shows the utility of Telomerality as an assembly quality Feature. Notably CHM13v2 shows a decreased QV score compared to previous version, however this is only due to the addition of the Y chromosome which comes from the HG002 cell line as was combined with CHM13 (Rhie et al. 2023). Without the Y chromosome, the average QV value of each chromosome was 75, comparable to the CHM13v1.1 calculated QV of 73.

For each run of paqman.sh we used varying numbers of threads (--threads) and calculated the run time and maximum RAM usage using *time* and *sacct* (--format=MaxRSS) respectively (Table 1; Table S1). For S288c using 16 threads, PAQman required on average 6minutes per assembly to complete and used a maximum of 5.5GB RAM. For Nipponbare, with a 31 times larger genome (∼12Mbp vs ∼380Mbp) PAQman required on average 93 minutes and a maximum of 48GB of RAM. For the human dataset CHM13 (∼3Gbp), we increased the thread count to 64 and saw that PAQman required on average 424 minutes and a maximum of 219 GB of RAM. In all datasets tested, speed improved with additional threads with minimal changes to max RAM usage. Using the Nipponbare assembly GCA000005425 and 16 threads, we isolated each major step in PAQman and saw that 65% of time and the highest RAM usage was during the steps containing read alignment with minimap2 and Correctness estimation with CRAQ (Table S2).

**Table 1:** Average run time of *paqman.sh* across five assemblies for each dataset using an AMD EPYC 7513 32-Core Processor and x86_64 architecture. Run time (real) was calculated using the *time* command in Linux; Max RAM usage was calculated using *sacct* (mean ± the standard deviation). Only 32 threads were tested on CHM13 with PacBio HiFi reads.

| Input |  |  |  |  | Run results |  |
| --- | --- | --- | --- | --- | --- | --- |
| Dataset | Genome size | Reads | coverage | threads | real (minutes) | Max RAM usage (GB) |
| S288c | ~12Mbp | ONT | ~100X | 8 | 6.92 ± 0.1 | 6.19 ± 0.78 |
| | | | | 16 | 6.29 ± 0.31 | 5.54 ± 0.36 |
| | | | | 32 | 5.3 ± 0.14 | 4.73 ± 0.42 |
| Nipponbare | ~380Mbp | HiFi | ~85X | 8 | 139.75 ± 1.96 | 42.88 ± 7.09 |
| | | | | 16 | 92.86 ± 1.71 | 47.864 ± 12.55 |
| | | | | 32 | 75.8 ± 2.66 | 47.31 ± 15.77 |
| CHM13 | ~3Gbp | HiFi | ~25X | 64 | 423.59 ± 23.32 | 218.67 ± 32.16 |

### Installation, Usage and Citation

PAQman was built for Linux operating systems. The easiest way to install PAQman is using the conda package manager:

> *conda config --append channels*
>
> *conda-forge conda config --append channels bioconda*
>
> *conda config --append channels pwwang*
>
> *conda install samtobam::paqman*

It may also be run as an Apptainer/Singularity image:

> *docker pull ghcr.io/samtobam/paqman:latest*

Both PAQman commands are run as below with default options (additional arguments available for paqman.sh):

> *paqman.sh -a path/to/assembly.fa -l path/to/longreads.fq*
>
> *paqplots.sh -s path/to/combined_summary_stats.tsv*

The PAQman GitHub (https://github.com/SAMtoBAM/PAQman) contains descriptions of installation/usage, commands, parameters, output files and both the features and their metrics.

PAQman has many dependencies that are stand-alone programs in their own right. Please cite both this manuscript in addition to its core dependencies. For example: “We used PAQman v1.2.0 (O’Donnell et al. 2025) in conjunction with Quast v5.3.0 (Mikheenko et al. 2023), BUSCO v6.0.0 (Tegenfeldt et al. 2025), meryl v1.3 (Miller et al. 2008), Mercury v1.3 (Rhie et al. 2020), Rasusa 2.2.2 (Hall 2022), CRAQ v1.0.9 (Li et al. 2023), BWA v0.7.19 (Li 2013), minimap2 v2.30 (Li 2018), samtools v1.22.1 (Danecek et al. 2021), mosdepth v0.3.12 (Pedersen and Quinlan 2018), bedtools v2.31.1 (Quinlan and Hall 2010), seqkit v2.10.0 (Shen et al. 2016) and ggplot2 v3.5.2 (Wickham 2016) to assess and visualize assembly quality.”

## Conclusion

As the ease of producing reference-quality assemblies continues to advance, new tools are needed to comprehensively evaluate assembly quality in a high-throughput manner. PAQman provides a simple means of evaluating an assembly’s overall quality by streamlining the calculation of seven commonly used features in the absence of a reference assembly. PAQman thus not only helps identify possible issues present within a specific assembly but simplifies the process of benchmarking assembly software to find the set of parameters that produce the “best” assembly for a given sequencing dataset and organism.

## Data Availability

PAQman is distributed as a conda environment and an Apptainer image. Source code and documentation are freely available through the GitHub repository at https://github.com/samtobam/paqman and a Zenodo archive at https://doi.org/10.5281/zenodo.16039705 for v1.2.0 used in this study.

## Author Contributions

Samuel O’Donnell (Conceptualization, Investigation, Methodology, Software, Validation,

Visualization, Writing – original draft, Writing – review & editing)

Ningxiao Li (Data curation, Writing – review & editing)

Jacob L. Steenwyk (Conceptualization, Writing – review & editing)

David M. Geiser (Resources, Writing – review & editing)

Frank N. Martin (Data curation, Resources, Writing – review & editing)

Emile Gluck-Thaler (Conceptualization, Funding acquisition, Resources, Writing – review & editing)

## Conflicts of Interest

JLS is an advisor to ForensisGroup Inc. JLS is a scientific consultant to Edison Scientific Inc.

## Funder Information

EGT and SO are supported by the Office of the Vice Chancellor for Research and Graduate Education at the University of Wisconsin-Madison with funding from the Wisconsin Alumni Research Foundation and the Department of Plant Pathology at the University of Wisconsin-Madison. JLS is a Howard Hughes Medical Institute Awardee of the Life Sciences Research Foundation.

## Notes

https://github.com/samtobam/paqman

https://doi.org/10.5281/zenodo.18175526

### Competing Interest Statement

JLS is an advisor to ForensisGroup Inc. JLS is a scientific consultant to FutureHouse Inc.

### Summary of Updates

Updated PAQman version from V1.1.0 to V1.2.0 Added additional benchmarking datasets

